# AlphaFold3 predicted LWO G-protein complex from European robin features active-state biased G_t_α

**DOI:** 10.64898/2026.05.19.726335

**Authors:** Jonathan Hungerland, Andrei Y. Kostritski, Karl-Wilhelm Koch, Ilia A. Solov’yov

## Abstract

Avian phototransduction and magnetoreception have been proposed to involve shared retinal proteins, including interactions between long-wavelength opsin (LWO), the cone-specific heterotrimeric G protein (G_t_), and cryptochrome 4a (Cry4a), yet structural information on avian phototransduction complexes is lacking. Here we present and critically assess two atomistic models of the European robin LWO–G_t_ complex generated by distinct modelling strategies. A full-complex prediction using AlphaFold3 yields a tightly packed, structurally stable interface but exhibits pronounced activation-like conformational features of the G_t_*α*-subunit that persist in simulations of the isolated protein, revealing a strong bias toward the active state. In contrast, a template-guided assembly based on single-chain predictions and an experimental rhodopsin-G_t_ reference structure forms a weaker interface and shows no intrinsic activation bias, while still displaying subtle activation-related dynamics. These results demonstrate that machine-learned complex prediction can encode functional states independently of the local interaction environment, thereby limiting its interpretability for signalling mechanisms that hinge on activation equilibria. Our findings highlight the need for explicit assessment of conformational-state bias when modelling regulatory protein assemblies and provide a structural framework for future studies of Cry4a-dependent modulation of retinal G-protein signalling in avian magnetoreception.

## Introduction

The primary steps of visual information processing in vertebrates take place in rod and cone photoreceptors. This phototransduction process has been studied intensely by physiological, biochemical and genetic approaches, but nearly all information has been obtained from studies on mammalian, amphibian, and fish species.^1,2^ Biomedical research, for example, employed very few vertebrate species as model organisms to understand hereditary human retinal diseases at the cellular and molecular level.^3–5^ In particular, many details of phototransduction in avian species remain unclear. Although similar genes mediate the molecular processes of bird phototransduction, the need for specific adaptations in diurnal or nocturnal species induces evolutionary pressure that has an effect at least on a genetic level.^6^ An example of a positively selected features within the phototransduction cascade is the faster recovery of the photoresponse, which is seen as a condition for better performance in motion detection and is an important sensory processing step for free flying birds.^7,8^ Futhermore, the absorbance of cone visual pigments varies among different bird species, probably reflecting a spectral fine-tuning due to specific habitat demands.^9,10^

Genetic adaptations to nocturnal activity of night-migratory songbirds might also have facilitated the use of the Earth’s magnetic field for seasonal navigation as the magnetic sense of night-migratory songbirds. Due to a wavelength-dependence of the magnetosensitive capabilities^11^ similar to photoexcitation of rod and cone cells, magnetoreception is likely also associated with processes in the retina. One of the currently discussed models to explain magnetoreception is the radical pair mechanism^12–15^ that is based on a photosensitive process in the retina. The involvement of the flavoprotein cryptochrome, particularly the isoform cryptochrome 4a (Cry4a), is further supported by observations on protein evolution,^16^ seasonal expression patterns^17^ and the effects of radio frequencies on navigation. ^18^ Cry4a is expressed in long-wavelength single cones and double cones of the European robin retina^17^ and exhibits magnetic-field dependent absorbance properties in response to blue light activation.^19^ The localization of Cry4a in cone cells matches results from a yeast-two-hybrid screening that aimed at identifying putative interaction partners of Cry4a.^8^ Among the identified candidates are the *α*– and *γ*-subunits (G_t_*α* and G_t_*γ*) of the cone specific heterotrimeric G protein, and the long wavelength-sensitive opsin (LWO).^8^ These results prompted further direct protein-protein interaction studies of Cry4a and G_t_*α* by surface plasmon resonance (SPR) spectroscopy, biochemical pulldown tests, and Förster resonance energy transfer measurements.^20,21^

Although the interaction of Cry4a and G_t_*α* is confirmed by different experimental approaches, a magnetic field sensitive signalling process in the bird retina is unknown so far. The two candidates G_t_*α* and LWO also form the trigger complex of the light-induced photo-transduction cascade. Thus, a possible interaction with Cry4a in cone cells raises questions about the structural organization of such a complex that might switch from a phototransduction to a magnetoreception state. A first study directed to compare quantitative binding data of G_t_*α* to either Cry4a or LWO was recently performed by Yee *et. al.*^21^ When peptides representing the cytoplasmic loops of European robin LWO were tested in interaction studies with G_t_*α*. The second loop connecting the transmembrane regions three and four showed high affinity binding^22^ pointing to a switching mechanism depending on ambient illumination. However, previous interaction studies lacked three-dimensional structural information of European robin specific proteins.

In the present study, we aim to achieve a structural view of the European robin G protein-LWO complex. We have used AlphaFold3 to predict an initial configuration of the protein complex and equilibrated it using all-atom molecular dynamics (AF3 model). To test this model, we have also prepared the same complex by leveraging the experimentally determined structure of the bovine LWO complex suggested by Gao *et. al.*^23^ and combined it with single chain prediction (Manual model). While the AF3 model showed a convincing, tight structural arrangement and activation-related dynamics, it also displayed crucial data biases. In particular, a clear bias towards the active state of the G-protein subunit was found. We ascribe this activation-related dynamics to a prediction related bias by recovering it in separate simulations of the extracted and solvated G_t_*α* subunit where the LWO was absent.

## Results

The LWO G-protein complex was constructed by two approaches. In the first approach, the protein complex was predicted using AlphaFold3 (AF3 model) and in the second, single chain protein structures were predicted and assembled (Manual model). The assembled AF3 complex after the first 1 *µ*s of simulation is shown in Fig. 1A. The LWO (cyan) is embedded in a lipid bilayer (gray surface) and contains all-trans-retinal. Under physiological conditions, 11-cis-retinal is photo-activated, which alters the electronic structure of the molecule such that the excited state of the retinal relaxes into the excited state minimum which is an all-trans-retinal configuration.^24,25^ After subsequent relaxation into the electronic ground state, a reverse rotation is energetically highly unfavourable because the required transition state features a broken *π*-bond.^26^ Thus, the intermolecular strain caused by the rotation resolves through the rearrangement of the LWO which most importantly includes the opening of a binding crevice.^27^ The AF3 model was predicted in a configuration with an open binding crevice. In the Manual model, the crevice was created during the optimization process with respect to the reference protein complex from *Bos Taurus* (pdb ID 6OY9^23^). Over all replica simulations of both models, LWO retained its active state configuration.

**Figure 1:**
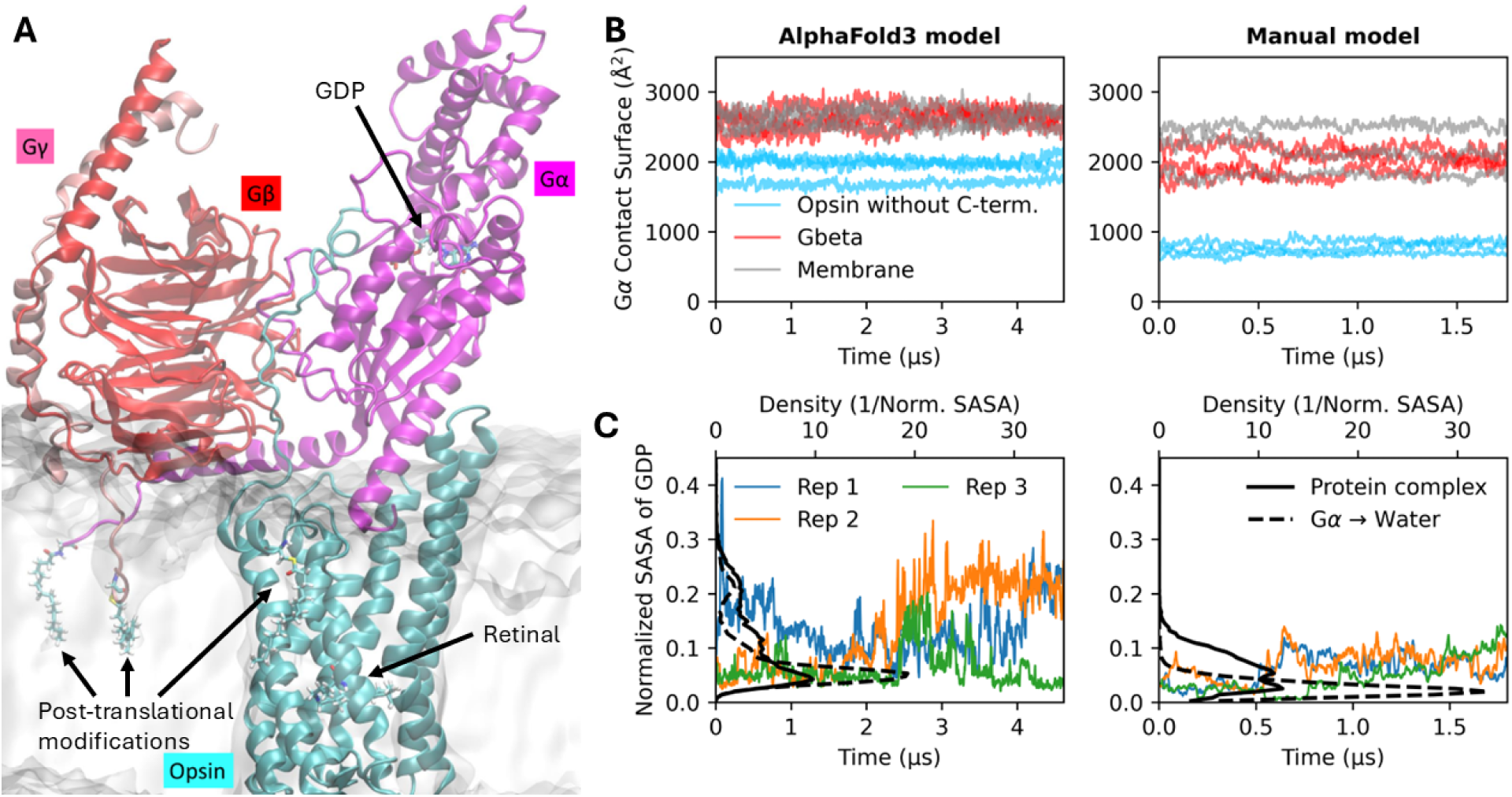
A: Rendering of the AlphaFold3 protein complex model after 1 *µ*s of equilbrium MD simulation (replica 2). The three-dimensional structure including secondary structural elements is shown for rhdopsin (cyan), G_t_*α* (magenta), G_t_*β* (red) and G_t_*γ* (light pink). Post-translational modifications (acylation of rhodopsin and G_t_*α*), the retinal unit and GDP are shown in Licorice representation and the membrane is indicated by a transparent, gray sur-face. **B:** Contact surface of the G*α* subunit with other system constituents over time. Right column: AF3 model, left column: Manual model. Replica simulations are not differentiated here. **C:** Normalized solvent accessible surface area (SASA) of the GDP in the protein com-plex simulations (colored lines) and the distribution of the data (solid black line). A black dashed line shows the GDP SASA distribution of separate simulations where the unequi-librated G*α* subunit was extracted from the protein complex model and placed in a water box.

An elongated, stretched out C-terminus of LWO was predicted for the AF3 model. In its initial configuration, the C-terminus traced upwards along the G_t_*α* subunit until it terminated close to the binding pocket of GDP. The C-terminus was kept in order to see whether the predicted but highly uncertain conformation was stable and whether it might play a role in G_t_*α* activation. The C-terminus eventually detached in all simulations without adopting its initial configuration again.

Post-translational modifications were included for both models. For the AF3 model, the usually myristoylated G_t_*α* subunit was already predicted with a membrane-embedded N-terminus, i.e. the N-terminus traced downwards approximately parallel to the opsin-helices. For all other post-translational modifications applied to the models, a dedicated refinement protocol was necessary to create reasonable starting configurations for the simulations. All 31 crystallized rhodopsin-G-protein complexes that we used for comparison did not feature post-translational modifications. Despite this fact, the orientation of the G_t_*α* N-terminus already indicated an embedding into a lipid membrane, in agreement with recent surface plasmon resonance experiments.^28^

A diverse range of LWO variants including those from mouse, chicken and zebrafish feature two palmitoylations at helix 8 which keep the helix close and orthogonal to the membrane.^29,30^ The sequence of european robin LWO used here features only a single cysteine in helix 8 such that only one palmitoylation was included. As expected, the palmitoyl chain interacts with the lipid environment and hydrophobic patches of the membrane-embedded rhodopsin. In all simulations, this helix remained stable and formed tight interactions with the lipid heads.

We evaluated all experimentally resolved protein complexes that feature G_t_*α* bound to rhodopsin. In all cases, the physiologically relevant binding of the G_t_*α* C-terminus into the activation-specific binding crevice of LWO was structurally conserved. Furthermore, aligning the 12 terminal residues of the considered 6OY9 template structure ^23^ with all other available complexes showed a high conservation of the terminus. To quantify the sequence similarities, a BLOSUM score matrix can be employed which expresses the likelihood of finding amino acid substitutions within conserved regions. In particular, aligning the 6OY9 terminus with any other position of the G_t_*α* sequenes yielded an average BLOSUM62 score of around 10 while the alignment at the C-terminal yielded a score of more than 60. Since BLOSUM62 scores measure sequence similarity on a logarithmic scale, the score difference indicates that the 6OY9 C-terminus sequence is orders of magnitude more likely to occur at the C-terminus than it is to occur at any other location of the sequence. Thus, a large number of similar complexes (31 structures) with high similarity at the key protein interface allowed the AF3 model to reproduce the expected binding of G_t_*α* to rhodopsin. Figure 1B shows the contact surface between G_t_*α* and the other protein subunits for the two studied models. The contact analysis for the AF3 model excluded residues 340 to 365 since the flexible C-terminus was not retained in the Manual model. The analysis shows that, across all simulations, the contact surfaces are relatively constant over time. However, G_t_*α* in the AF3 model (left column) shows a higher contact area to the opsin, to G_t_*β* and to the membrane than the Manual model. This observation highlights the tight arrangement of the AF3 predicted complex. In contrast, the Manual model shows only comparably shallow contacts to opsin. The pursued modelling ansatz for the Manual model that replicates a gradual approach of the single protein structures initially only allows interfacial contacts. In order to achieve a tighter, interlocked arrangement, atomic repulsions should have been turned off in the early stages of the complex relaxation. Similarly to the gas-particle-like initialization of atomic nodes for diffusion-based structure prediction models like AF3,^31^ an initial neglect of atomic repulsions would ease configurational energy barriers towards the energetic minimum of the complex structure. Within the context of the physical realism of the model, the Manual model should be considered to represent the early stages of the complex formation. The AF3 model on the other hand, is representative for a later stage of the protein complex, as discussed below.

The *α*-subunits of heterotrimeric G proteins contain in general two relevant interaction domains: the RAS-homology domain and the *α*-helical domain.^32^ While the RAS-homology domain of G_t_*α* interacts with rhodopsin, the *α*-helical domain sits on top of the GDP, preventing its unbinding. The interaction between LWO and G_t_*α* causes the *α*-helical domain to open up extensively such that G_t_*α* adopts its active conformation in which GDP to GTP conversion can occur similar to changes observed in G_s_-protein in complex with the *β*_2_-adrenergic receptor.^32^ Even though GDP never left its binding pocket in the course of the simulations, the solvent accessible surface area (SASA) of GDP is a suitable measure to inform on the activation of G_t_*α* because an increased SASA can be expected to correlate with reduced GDP binding affinity as well as a reduced unbinding energy barrier. The SASA was calculated in VMD^33^ such that the G_t_*α*-GDP complex surface was restricted to sampling points around the GDP and it was normalized by the surface area of GDP in its current configuration if it had been surrounded purely by water. For the AF3 model, the normalized SASA with a running window average of 10 ns is shown over time in Fig. 1C (left, colored lines). The first replica simulation features a high jump in SASA up to 40% and then relaxes until it rises again at around 4 *µ*s. For replica 2, there is a more steady and stable increase after 2 *µ*s of simulation while replica 3 only shows an intermediate rise in SASA. Replica simulation 2 indicates that the model structure is realistic in the sense that it can exhibit the physiologically relevant interactions and can reproduce the mechanical interactions that lead to G_t_*α* activation. Replica 1 on the other hand suggests that the AF3 model may be subject to data bias towards the activate state since the mechanical interactions leading to the activation process would naturally not occur on timescales of a few nanoseconds during which the initial peak in SASA is formed. The GDP SASA of the Manual model (see Fig. 1C, right, colored lines) remains at a value of about 0.05 until it increases slightly after 0.5 *µ*s for replicas 1 and 2, while replica 3 shows a complete closer after the same amount of simulation time.

To investigate the potential bias on the G_t_*α* conformation further, additional simulations were performed in which the G_t_*α* subunit from both initial models were extracted and placed into a waterbox with 0.15 mol/L KCl. The extracted G_t_*α* units were simulated for 8 replicas over 800 ns each. Figure 1D compares the probability density of GDP SASA for the complex simulations and the solvated G_t_*α* subunits. The Manual G_t_*α* subunit placed in water exhibits a single clear peak at 0.03 normalized SASA. In the Manual complex model, there is a small but significant population at increased SASA of 0.1. This result suggests that the Manual complex model captures the mechanical interactions that induce G_t_*α* activation despite its rather interfacial interaction. For the AF3 G_t_*α* protein subunit on the other hand, there is an increased SASA of the closed conformation at 0.08 and a small population with over 0.2 normalized SASA. Comparing the simulations of G_t_*α* in its protein complex and in water, shows that the conformations of highly increased SASA at over 0.2 exist in both sets of trajectories. Thus, the G_t_*α* protein structure as predicted by AlphaFold3 in its complex conformation already includes a conformational bias towards the active G_t_*α* state. For the AF3 predicted protein structure, the LWO may not even be necessary to induce complete G_t_*α* activation.

To gather more information on the exact parts of G_t_*α* that are subject to activation-related sturctural differences, we analyzed the fluctuations and inter-protein contacts as shown in Fig. 2A. The average root-mean-square fluctuation of the backbone atoms of G_t_*α* are shown for each residue, while colored bands highlight contacts to other proteins. The *α*-helical domain including residues 61-170 rests upon the RAS domain such that the *α*-helical domain can open on one side like the top to a chest with a hinge. Residues further down the sequence are in close contact with the rhodopsin, G_t_*β* and G_t_*γ* subunits. Some of those contacts coincide with increased dynamic fluctuations of the backbone atoms.

**Figure 2:**
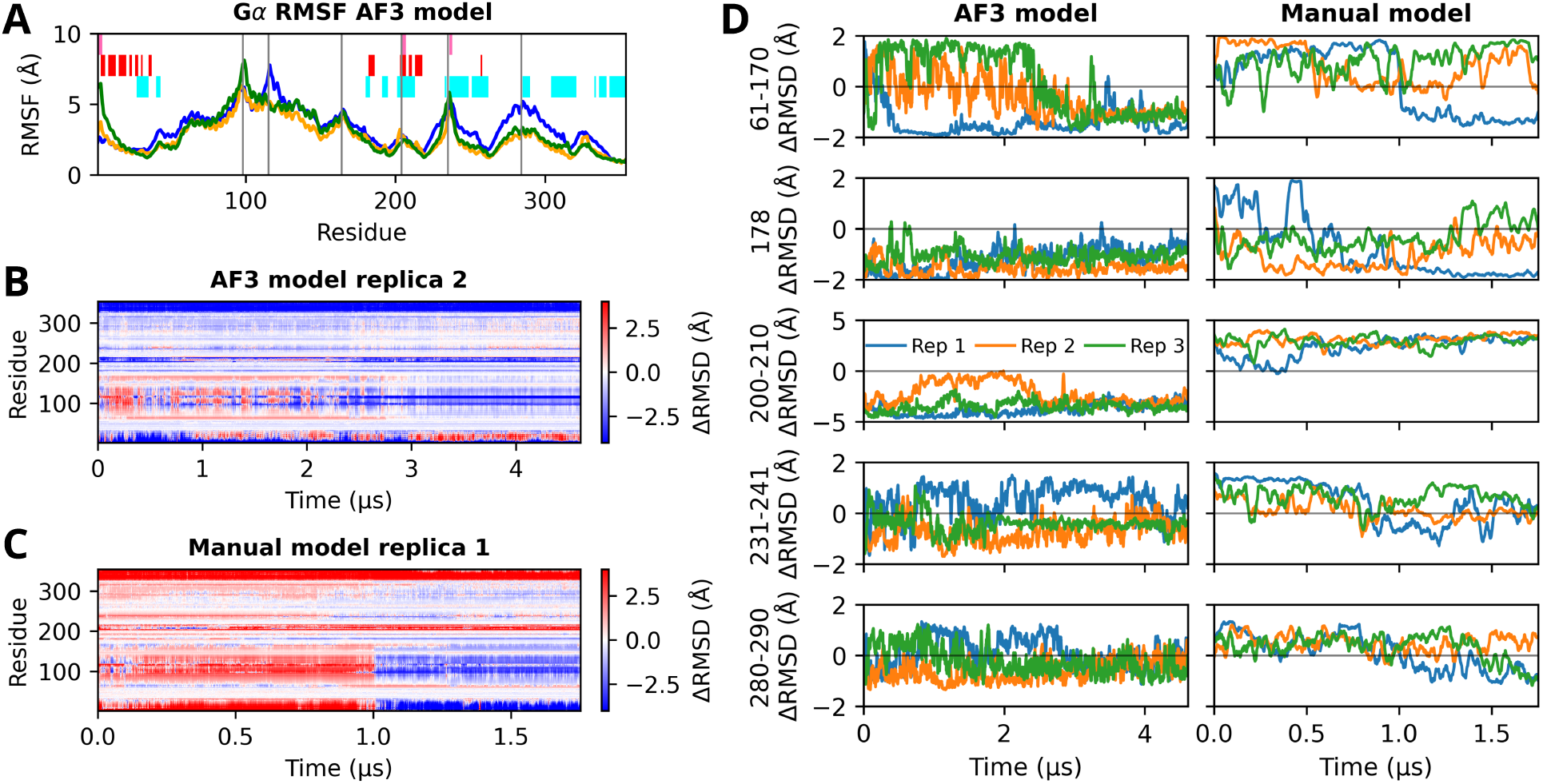
A: Root-mean-square-fluctuation of the backbone atoms of the G_t_*α* subunit for the replica simulations 1 (blue), 2 (orange) and 3 (green) of the AF3 model. Colored bands indicate stable contacts (within 5 Å over at least 95% of the simulation time) to LWO (cyan), G_t_*β* (red) or G_t_*γ* (light pink). The peaks indicated by grey vertical lines are located at residues 98, 115, 164, 204, 235 and 284. **B:** Backbone atoms ΔRMSD as defined by Eq. (1) for replica simulation 2 of the AF3 model separated by residues and timepoint. The colorscale is capped at ±4 Å. **C:** Same as B but for replica simulation 1 of the Manual model. **D:** ΔRMSD averaged over different selections of residues. Results for replica simulations are color-coded in blue (1), orange (2) and green (3).

To further characterize the current configuration of each protein residue, we have introduced and computed the ΔRSMD value that is defined as follows:

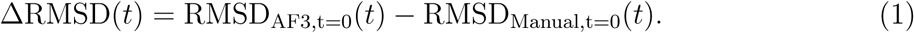

Here, the RMSD indicates the root-mean-square deviation across the G_t_*α* backbone atoms of the protein residue after aligning the G_t_*α* protein structure using the C-*α* atoms of residues 35 to 40, 46 to 55, 195 to 200, 220 to 225 and 264 to 268 as reference positions. For RMSD_AF3,t=0_, the initial, unequilibrated AF3 model was used for alignment, while in the case of RMSD_Manual,t=0_, the Manual model was employed. Positive values of the ΔRMSD show that the residue conformation is closer to the Manual initial model than to the AF3 initial model. For negative values, the residue conformation is closer to the AF3 initial model.

Figure 2B shows the per-residue values of the ΔRMSD over time for replica 2 of the AF3 model. A thick, dark blue band at the C-terminal residues illustrates that the conformation of the C-terminus appears to be at least 4 Å closer to the AF3 model throughout the entire simulation time. A pattern that was found for all replica simulations and is similarly consistent for the Manual models as shown in Fig. 2C where the C-terminus configuration is closer to the initial Manual model instead.

A noticeably different pattern occurs across the *α*-helical domain (residues 61-170). Replica 2 of the AF3 model initially switches between AF3-like and Manual-like conformations until it falls into and remains in an AF3-like conformation after 2.5 *µ*s – the same at which the GDP SASA (see Fig. 1C) was found to increase. The Manual model replica 1 simulation, on the other hand experiences a clear shift towards a configuration closer to the AF3 model at 1 *µ*s.

A more encompassing analysis is given in Fig. 2D, where the averages over selected regions of the ΔRMSD values are shown. Starting with the *α*-helical domain of residues 61-170, it can be noted that all replica simulations of the AF3 model eventually adopt an AF3-like conformation after 2 *µ*s. This finding underscores the realism of the initial prediction but it also supports the observation of an active state bias where the *α*-helical domain is already initialized in a configuration that tends towards GDP release. In the Manual trajectories, the *α*-helical domain transitions into an AF3-like conformation. While *α*-helical domain of all three Manual replica simulations explores conformations between the Manual initial model and the AF3 initial model, replica simulation 1 in particular features a clear, nearly step-wise configurational change after about 1 *µ*s of simulation time.

The ΔRMSD of the key switch residue ARG178 from the G_t_*α* subunit is plotted in the second row of Fig. 2D. For all AF3 model trajectories, Manual-model like configurations are only very short-lived intermediates, while the Manual-model trajectories show a transition towards an AF3-like configuration that is either rapid (replicas 2 and 3) or gradual but highly consistent (replica 1). Interestingly, the configurational switch of ARG178 in Manual replicas 1 and 2 clearly precedes the change in the *α*-helical domain, which supports a possible causality. Residue ARG178 corresponds to the so-called arginine finger that controls GTPase activity.^34,35^ In the inactive state, it binds to the phosphate groups of GDP. The disruption of this salt bridge is a key step in G-protein activation (residue ARG201 in^32^). In the range of residues 200-210, the two modelling approaches exert two clearly separate configurations. At this location, G_t_*α* is in contact with rhodospin, G_t_*β* and G_t_*γ* (see Fig. 2A), making it a key contact region. While the ΔRMSD of residues 200-210 suggests that the configurational differences in this region remain constant over time, residues 231-241 and 280-290 are more dynamic and in fact the Manual model appears to adopt the AF3 conformation over time. The residues 280-290 of the AF3 model trajectories retain an AF3-like conformation and the Manual models adopt comparable structures after about 1 *µ*s. As this region is expected to partake in the mechanical interaction leading to G_t_*α* activation, the G_t_*α* C-terminal ΔRMSD suggests that the AF3 model contains conformational insights that would otherwise only be found after microseconds of molecular dynamics simulations.

Figure 3 directly overlays the initial model structures. The overall folds of the protein complexes displayed in the centre of the image are similar. Noteworthy differences of the AF3-model are the deep intrusion of the G_t_*α* C-terminus into the Opsin binding pocket, the G_t_*α* N-terminus embedding into the membrane patch and the G_t_*α α*-helical domain tilting to the left.

**Figure 3:**
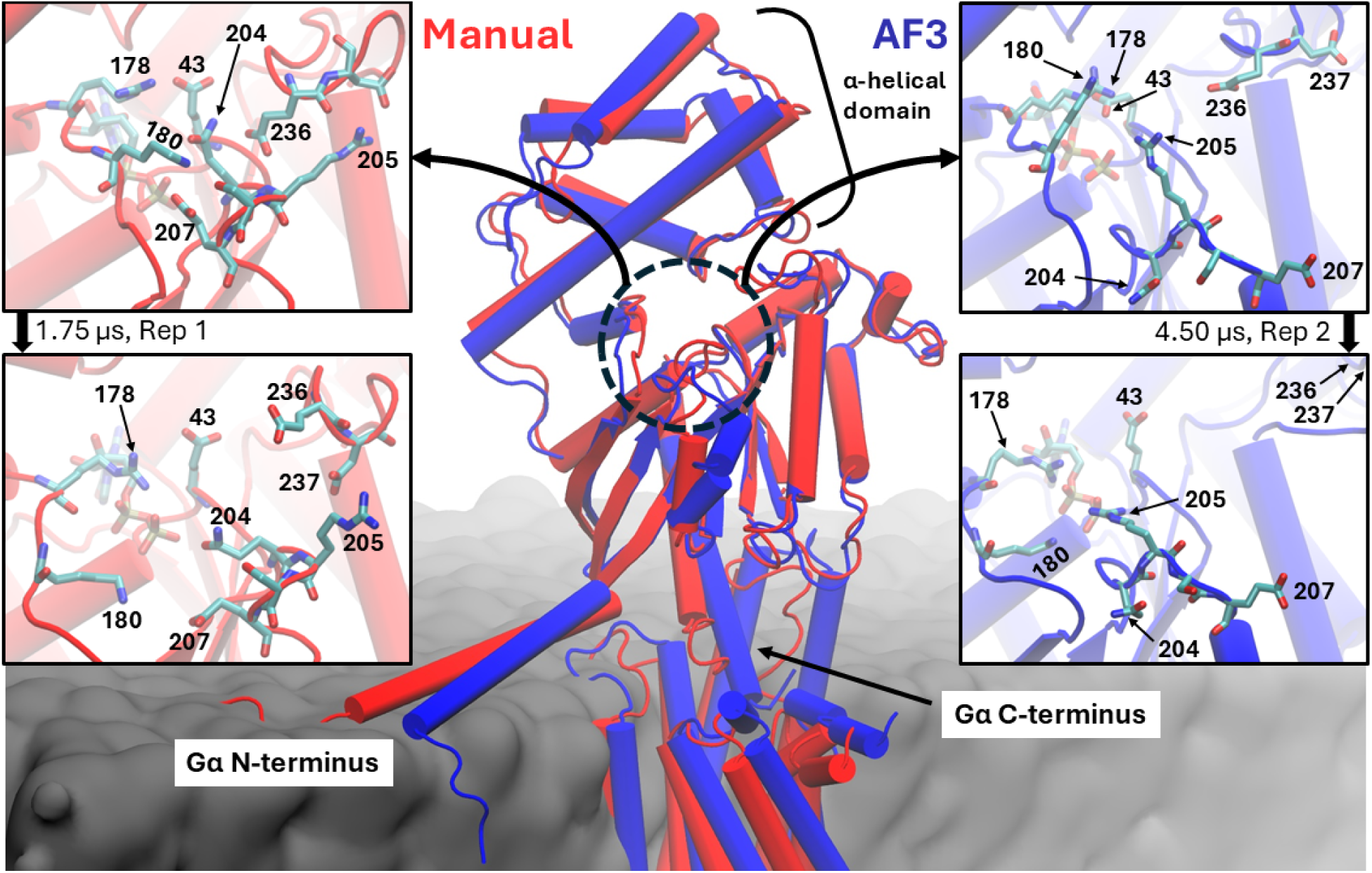
Comparison of the initial model structures. LWO and G_t_*α* from the Manual (red) and AF3 model (blue) are overlayed, a cut through the membrane surface (grey) is shown for orientation. Left and right: zoom-in on the region that appeared in ΔRMSD analysis. Top panels: initial models before molecular dynamics, bottom panels: the structure of selected replicas after the indicated simulation time. Hydrogens were omitted for clarity.

A closer look on the sidechain of residues 200-210 reveals multiple ionic and hydrogen bonds in the initial Manual model that protect the GDP binding pocket. An electrostatically stabilized network of residues 178, 43, 204, 180 and 207 sits on top of the phosphate groups of GDP. In replica simulation 1 of the Manual model, which features a characteristic change into an AF3-like conformation after 1 *µ*s, the electrostatic interaction network is disturbed beyond recognition. While the salt bridge betwen LYS180 and GLU207 is not entirely disrupted, the involved residues are heavily displaced, such that the GDP becomes solvent accessible. At the end of the simulation, ARG178 exerts pronounced interactions with the phosphate groups of the GDP, supporting its central role in G_t_*α* functionality.^32,34,35^

The AF3 initial model structure provides much weaker protection for the GDP binding pocket. The tight electrostatic network observed in the Manual model is already dispersed. After 4.5 *µ*s of simulation, the loop containing acidic residues 236 and 237 opens outwards, allowing water molecules to penetrate a key interaction site between the *α*-helical domain and the Ras-domain, which is partially responsible for the increased GDP solvent exposure. ARG178 and ARG205 rotate towards the GDP, which retains a favorable GDP binding pocket at the expense of further weakened *α*-helical-domain and Ras-domain interactions.

The *α*-*β* loop of residues 200-210 distinguishes experimentally resolved G_t_*α* structures that were associated in a GPCR complex. Figure 4 displays the loop for the two models and four crystal structures. The previously discussed electrostatic interactions of pure G_t_*α* and their absence in complex-associated G_t_*α* structures can both be seen in the crystal structures. Furthermore, the *α*-helical part of the *α*-*β* is partially unfolded in the complex-associated G_t_*α* structures as well as in the AF3 model. Note that the *α*-helical domains of the chosen G-protein complex crystal structures are either unresolved (6OY9) or completely flipped open (6PT0), such that there are no contacts between the *α*-helical domain and the Ras domain of the G-proteins. The Ras-domain structure of the AF3 model mimics that of the complex-associated crystal structures, even though the *α*-helical domain is still in the closed conformation in the initial AF3 model. The AF3 prediction is therefore a computational chimera between a closed G_t_*α* model and an activated G_t_*α*.

**Figure 4:**
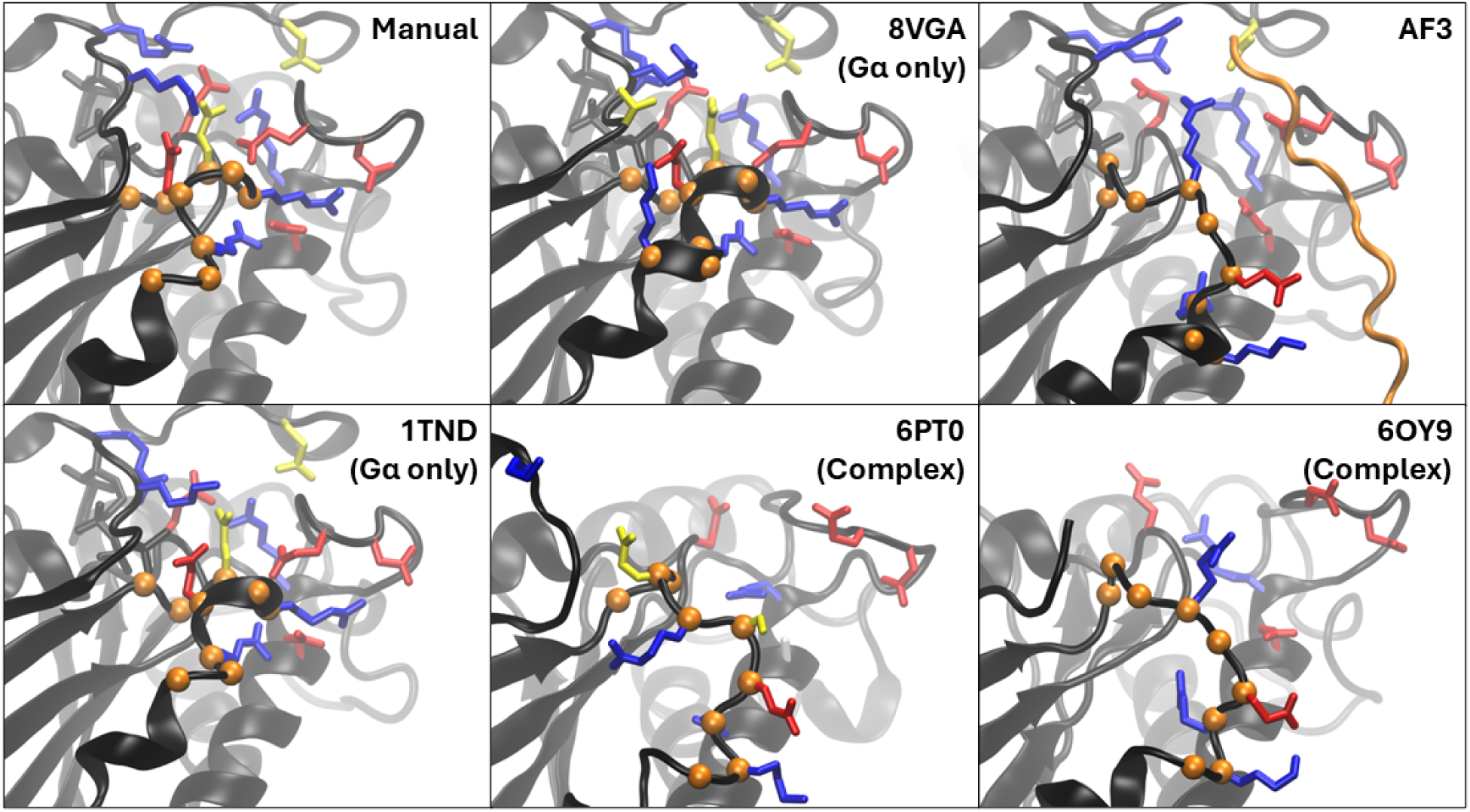
Comparison of the GDP-proximal loop region that differentiates the Manual and AF3 models for different crystal structures as indicated by their pdb IDs. Black: G_t_*α* subunit and GDP ligand. (Blue/Red/Yellow): Selected amino acids that are (positively charged/negatively charged/neutral polar). Orange spheres: C*α* atoms of residues 202-210 in the *European* robin G_t_*α* sequence and their corresponding locations after alignment for other structures. Orange line: rhodopsin C-terminal in the AF3 model.

The direct comparison between experimentally resolved complex models with the AF3 predicted structure puts the location of the G-protein coupled receptor C-terminal tail into a new perspective. The C-terminal disrupts the electrostatic network which protects the GDP binding pocket in the Manual model and it competes with the GDP phosphate groups for salt bridges with positively charged residues. Even though the receptor C-terminus is often unresolved, some studies measured parts of it tracing upwards along a cleft between the *α*– and *β*-subunits,^36,37^ which agrees with the prediction in the AF3 model. Even though G-protein activation does not entirely deplete upon truncation of the receptor C-terminus, the C-terminus was found to mediate G-protein specificity.^38,39^

## Dicussion and Conclusions

In this manuscript, we have presented two models for the *European robin* LWO G-protein complex. The first model was predicted with AlphaFold3 and the second model combined single protein prediction with a classical mechanics protocol that used information from an experimental template structure to model the protein interface. Our simulations have demonstrated a high quality of the AF3 model due to its ability to predict a highly stable protein complex with closely intertwined protein interactions that would otherwise be quite challenging to represent. However, we could establish that the AF3 model also brings along a crucial bias towards the G_t_*α* activation. Drawing on various experimental data that partially includes protein complexes in their activate state, the model already shows conformational features in the *α*-helical domain, residues 231-241 and residue 178 which would otherwise be induced by interactions at the protein interface. In our second protein complex model, the potential data bias was both limited to the protein interface and to a single, known template structure as the potential source of artifacts. While the model exerted a more shallow interface, it did not suffer from activation-state bias and yet still showed signs of activation-related conformational changes as indicated by a systematic, slight increase in GDP SASA, the conformational dynamics of residue 178 and the *α*-helical domain.

For the purpose of exploring functional protein complexes, our findings stress the importance of treating proteins as dynamic soft-matter systems. While static structures are informative, their functionality ultimately emerges from the formation of transient signalling assemblies that are embedded in a spatially and temporally organized interaction network.^40^ Crystallography or cryo-EM allows structural insights into this network. Machine learning methods based on experimental structures may construct a model that mimics measurement results and can adopt both biases and hidden insights. For the particular case of G-protein activation, we conclude that the presence of an expected conformational change in molecular dynamics simulations of the machine learned prediction is not sufficient to proclaim a realistic modelling of the environment interactions. Hidden data biases are possibly adopted from the experiment into the prediction, which must be accounted for when utilizing the models. However, the ability to reproduce the data biases also underscores that the machine learning model parameters contain information of functionally relevant conformational changes. In fact, it is possible to fine-tune protein structure predictions towards the reproduction of an ensemble of structures along a key transition coordinate.^41^

Within the context of potential interactions between the LWO complex and Cry4a,^8,20^ the observed data bias becomes even more relevant. Their combination would constitute a highly novel protein complex, given that cryptochrome 4 from any species has not been structurally resolved in contact with a second protein and that avian cryptochrome itself thus far is only resolved for a single species.^42^ Nevertheless, the previously observed interactions between Cry4a and proteins that partake in visual signal transduction^8,20^ may hint at a possible involvement of the LWO complex in the transduction of magnetic field information. In particular, it can be imagined that a protein-complex formation of e.g. G_t_*α* with Cry4a may depend on the activation state of Cry4a and that this complex formation may down-or upregulate G-protein activation. To test this hypothesis, one may build and study such protein complexes *in silico*. However, any protein complex prediction with an inherent bias on the activation process of G_t_*α* would skew these results.

To circumvent problems with the inherent bias described above, one may instead rely on less sensitive observables like the possible formation of a Cry4a-G_t_*α* complex that sterically prevents the association of G_t_*α* with rhodopsin, leading to downregulation of the G_t_*α* activity. Alternatively, interactions of Cry4 with the *α*-helical domain of fully activate G_t_*α* could effectively lock G_t_*α* into the activated state, leading to upregulation. As an additional boundary condition, the formation of functionally relevant complexes must also be dependent on the activation state of Cry4a. Otherwise, the complex formation would not represent any information about the magnetic field.

Nevertheless, the AF3 model may encode biologically and functionally relevant information. Even though the rhodopsin C-terminus was predicted with low confidence, it provides a reasonable initial structure to study G-protein specificity^38^ in the future. At least on a qualitative level, the AF3 predicted rhodopsin C-terminus supports the relevance of such a flexible region for G protein activation. However, to conclusively evaluate the mechanism of G-protein recognition,^39^ future computational modelling approaches must go beyond static structures and beyond the limits of atomistic molecular dynamics.

## Methods

### AlphaFold3 complex model

A rhodopsin-G-protein complex was predicted by AlphaFold3^31^ using amino acid sequences from the European Robin long-wavelength-opsin^22^ and the G-protein subunits *α*,^8^ *β*^43^ and *γ*.^44^

The protein complex was then embedded in a lipid membrane that replicates the composition that was determined for cone-photoreceptor cells of chicken.^45^ For the generation of the membrane and its combination with the LWO subunit, the CHARMM-GUI^46^ webserver and its membrane builder^47,48^ were employed. The G-protein subunits, the GDP and the retinal were then merged with the membrane-embedded rhodopsin. A covalent bond between the LYS312 residue of LWO and retinal was modelled via a Schiff-base with the retinal being fixed in the all-trans configuration characteristic for the structure after light-activation. ^24,25^ Other opsins were found to have palmitoylations,^29^ such that the CYS332 residue in helix 8 of LWO was modelled to be palmitoylated. CYS126 and CYS203 residues of LWO formed a disulfide bond that connects the third transmembrane helix and the second extracellular loop. The G_t_*α*-subunit received the lipid-like myristoylation at the glycine residue located at its N-terminus. Towards the N-terminal residue LEU68 of the G_t_*γ* unit, residue CYS65 was farnesylated. Following the protonation state prediction of Propka3,^49^ the ASP99 residue of LWO was protonated. System preparation was done using VMD.^33^

The constructed protein-membrane-complex was then subjected to atomsitic molecular dynamics (MD) simulations. All MD simulations were performed using NAMD3^50,51^ and employed the CHARMM all-atom force-field.^52–55^ Simulations were streamlined through the NAMD AutoConf package provided alongside all input and configuration files. A summary of the employed simulations protocols is given in the following. Unless otherwise stated, the default settings generally recommended for NAMD3 in its manual were used.

While the myristoylated N-terminus of G_t_*α* already showed a reasonable embedding into the membrane, the initial orientations of the farnesylation and the palmitoylation did not reflect their expected role in anchoring the G_t_*γ* C-terminus and the membrane-orthogonal helix 8 of rhodopsin. Therefore, a combination of free-energy perturbation, harmonic restraints and collective variables were used. Atoms within a radius of less than 15 Å around the G_t_*γ* farnesilation and the LWO palmitoylation were not restrained. In a radius between 15 Å and 30 Å, restraints were linearly introduced with force constants to 10 kcal/mol at a 30 Å distance. For all atoms further away, the force constant was set to 10 kcal/mol/Å.

To allow free movement of the initially misplaced farnesylation and palmitoylation, their non-bonded interactions were completely decoupled from the rest of the system via the al-chemical method. Simultaneously, harmonic forces were employd to pull the modifications into a conformation that points straight down into the membrane. These restraints were linearly increased from 0 kcal/mol/Å to 0.5 kcal/mol/Å over 1000 steps with a 0.25 fs timestep. During a second stage of 10k steps, non-bonded interactions between the modifications and the environment were recovered via a the alchemical *λ* parameter while the restraints were again increased from 0 kcal/mol/Å to 0.5 kcal/molÅ. With the fully interacting system in place, the harmonic forces to pull the modifications downward were further increased from 0.5 kcal/mol/Å to 5 kcal/mol/Å over 90k steps. Free energy changes of less than 1 kcal/mol, a final free energy difference of –3.6 kcal/mol and visual inspection confirmed that this approach led to a reasonable, membrane-embedded configuration. In a final gradient descent minimization for 20k steps, the harmonic forces that pulled the modifications downward were linearly removed over the first 10k steps.

The equilibration of the full system was initiated with 30k conjugate gradient minimization steps. An initial NVT simulation with a 0.1 fs timestep, increased radius of 15 Å for explicit (non-PME) non-bonded interactions and a Langevin damping of 1 ps^-1^ was performed for 10 ps during which everything except water atoms was restrained. For the next stage, the timestep was increased to 1 fs and the pressure control was employed at 1 atm. In this simulation state, the restrains were lifted on the membrane atoms except for oxygen and nitrogen atoms (which fixed the head groups) together with the restraints on residues 1-9 of G_t_*α* and 61-65 of G_t_*γ*; those settings were employed for 1 ns of simulation time. The non-bonded interactions were then switched to the default values (12 Å cutoff and 10 Å switchdist), the Langeving damping constant was increased to 5 ps^-1^ and full electrostatics were only computed every second simulation step. With only the backbone atoms of the protein complex (except residues 1-9 of G_t_*α*and 61-65 of G_t_*γ*) restrained, the simulation was continued for another 5 ns. The simulation was extended by another 5 ns, where all constraints were lifted and the simulation timestep was increased to 2 fs. Finally, the simulation timestep was increased to 4 fs (possible through hydrogen mass repartitioning^56^) and the NPT simulation was run for 4.5 *µ*s. Three replicas from identical initial coordinates were simulated with the above described protocol way.

### Manual prediction combined into complex

As an alternative aproach, the protein complex was generated by combining predictions of each protein subunit into a complex. The separate protein chains were predicted using the OpenFold^57^ implementation of the AlphaFold2^58^ algorithm. All crystallized protein complexes that contained a G-protein coupled receptor together with all G-protein subunits (pdb IDs 6CMO, 6D9H, 6G79, 6LMK, 6LML, 6NBH, 6OY9, 6OYA, 6PTO, 6QNO, 7DFL, 7DH5, 7FIN, 7FIY, 7JJO, 7S0F, 7S0G, 7T8X, 7T94, 7T96, 7UM5, 7UM6, 7UM7, 7X8R, 7X8S, 7XJH, 7XJI, 7Y12, 7Y15, 7YS6, 8SAI) were used to prepare candidate complexes. Sequence alignments were done using STAMP^59^ as implemented in the MultiSeq^60^ tool of VMD.^33^

To generate protein complexes, the predicted protein chains were first aligned with their respective counterparts of the template crystal structure. The template-aligned target complex was then prepared for a constrained minimization MD simulation. For all residues, where the sequence alignment did not result in a gap in the template structure sequence or the European robin sequence, the backbone atoms were restrained to overlay. To apply the restraints locally to the protein-protein interface, the interface weight function *f* (*d*) was defined as:

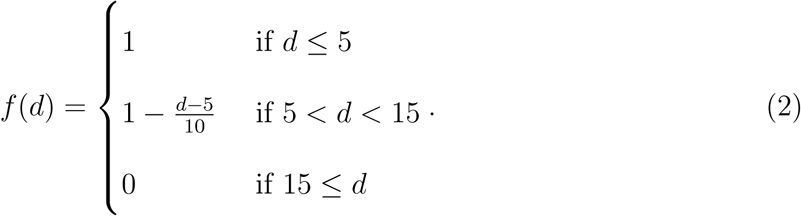

For two protein chains A and B that form an interface, the distance *d* was calculated for all residues of both structures as the minimal distance to any atom of the other chain. To compute the interface weight per residue, the reference crystal structure was used.

To allow the protein interfaces to relax, the previously aligned protein complex that consisted of the single protein predictions, were shifted away from each other by 9 Å. In particular, LWO was shifted away from G_t_*α*, while G_t_*β* and G_t_*γ* were shifted away from G_t_*α*. G_t_*γ* is additionally shifted away from G_t_*β*. The shifts were executed sequentially in the described order and the optimal direction to ensure a roughly orthogonal vector with respect to the interface which was determined visually. The shift distance was held constant via positional restraints on the atoms with respect to a shifted reference complex. The restraint force constants were initialized at 32 kcal/mol/Å and were weighted by *f* (*d*). Every 5k conjugate gradient minimization steps, the shift distance was reduced by 1 Å and the restraint force constant was halved until the shift distance reached zero. At zero shift distance, the restraint force constant was reinitialized at 32 kcal/mol/Å and again halved after every 5k minimization steps over a total of 50k minimization steps.

The described minimization protocol was performed on all generated protein complexes to find the most viable target for modelling. The sequence alignments genereated during the initial structure alignment were used to compute total BLOSUM scores and interface-weighted BLOSUM scores using the weight function defined in Eq. (2). The root-mean-square deviation (RMSD) was computed between all minimized complexes. The RMSD was used to cluster the complex candidate structures. The complexes generated from the templates 6PTO,^61^ 6CMO^62^ and 6OY9^23^ had an RMSD of less than 2 Å with each other and together formed the cluster with the highest average BLOSUM and interface-weighted BLOSUM score. From those three most promising complexes, 6OY9 was selected to proceed to MD simulation.

The subsequent preparation of the system was identical to that of the AlphaFold3 complex with the exception that the AlphaFold3 complex had a larger surrounding membrane patch. This was done because the initially simulated Manual complex model seemed to induce a potentially relevant curvature on the lipid environment. The initial equilibration phase was identical to the AlphaFold3 complex, described earlier. For the Manual complex, however, the first unrestrained stage was simulated for 50 ns at a 1 fs timestep, which was followed by 700 ns at a 2 fs timestep. Finally, another 1000 ns long isochoric simulation was performed that employed a 4 fs timestep.

## Acknowledgement(s)

IAS thanks the Deutsche Forschungsgemeinschaft (DFG) for its support through TRR386, HYP*MOL, project number 514664767; IAS, KK, and JH thank the DFG for funding through SFB 1372, Magnetoreception and Navigation in Vertebrates, No. 395940726 and EXC-3051, Excellence Cluster NaviSense, No. 533653176. IAS is grateful to the Ministry of Science and Culture of Lower Saxony (Dynamik auf der Nanoskala: Von kohärenten Elementarprozessen zur Funktionalität (DyNano)). AYK thanks DFG for funding through the Walter Benjamin programme, project no. 541484920. The authors gratefully acknowledge the computing time granted by the Resource Allocation Board and provided on the super-computer Emmy/Grete at NHR-Nord@Göttingen as part of the NHR infrastructure. The calculations for this research were conducted with computing resources under the project nip00058.

## Author contributions

JH and KWK wrote the initial draft. JH produced simulations, analysis and visualizations. JH and AK interpreted the data. IAS and KWK provided funding and resources. IAS, KWK and JH conceptualized the study. All authors were involved in scientific discussions. All authors reviewed the manuscript.

